# Multimodal Hippocampal Subfield Grading For Alzheimer’s Disease Classification

**DOI:** 10.1101/293126

**Authors:** Kilian Hett, Vinh-Thong Ta, Gwenaëlle Catheline, Thomas Tourdias, José V. Manjón, Pierrick Coupé, Alzheimer’s Disease Neuroimaging Initiative

## Abstract

Numerous studies have proposed biomarkers based on magnetic resonance imaging (MRI) to detect and predict the risk of evolution toward Alzheimer’s disease (AD). While anatomical MRI captures structural alterations, studies demonstrated the ability of diffusion MRI to capture microstructural modifications at an earlier stage. Several methods have focused on hippocampus structure to detect AD. To date, the patch-based grading framework provides the best biomarker based on the hippocampus. However, this structure is complex since the hippocampus is divided into several heterogeneous subfields not equally impacted by AD. Former *in-vivo* imaging studies only investigated structural alterations of these subfields using volumetric measurements and microstructural modifications with mean diffusivity measurements. The aim of our work is to study the efficiency of hippocampal subfields compared to the whole hippocampus structure with a multimodal patch-based framework that enables to capture subtler structural and microstructural alterations. To this end, we analyze the significance of the different hippocampal subfields for AD diagnosis and prognosis with volumetric, diffusivity measurements and a novel multimodal patch-based grading framework that combines structural and diffusion MRI. The experiments conducted in this work showed that the whole hippocampus provides the most discriminant biomarkers for advanced AD detection while biomarkers applied into subiculum obtain the best results for AD prediction, improving by 2% the accuracy compared to the whole hippocampus.

## Introduction

Alzheimer’s disease (AD) is an irreversible neurodegenerative process leading to mental dysfunctions. Subjects presenting Mild Cognitive Impairment (MCI) have higher risk to develop AD^1^. To study the preclinical phase of the disease, the Alzheimer’s disease neuroimaging initiative (ADNI) was set up based on two MCI definitions: early MCI (eMCI) and late MCI (lMCI). eMCI represents subjects with cognitive impairment milder than lMCI which is composed of amnesic MCI^2^. Such clinical symptoms caused by changes like synaptic and neuronal losses that lead to structural and micro-structural alterations. Neuroimaging studies performed on AD subjects revealed that brain structure alterations are advanced when diagnosis is established and emphasized the need to study the early stages of the disease.

The improvement of medical imaging techniques such as magnetic resonance imaging (MRI) enabled the development of efficient biomarkers detecting alterations caused by AD^3^. Over the past years, many methods have been proposed to perform automatic detection of alterations associated with AD. First, studies proposed methods based on specific regions of interest (ROI) capturing alterations at an anatomical scale. Among structures impacted by AD, previous investigations focused on hippocampus (HIPP)^4–6^, entorhinal cortex (EC)^7–9^, parahippocampal gyrus, amygdala^10^ or parietal lobe^11, 12^. Alterations on these structures are usually estimated using volume^13, 14^, shape^15, 16^, or cortical thickness^17, 18^ measurements. Second, beside ROI-based methods, whole brain analysis performed on structural MRI (s-MRI) have been proposed to detect areas impacted by AD at a voxel scale. These methods are usually based on voxel-based morphometry (VBM) or tensor-based morphometry (TBM) frameworks^19^. It is interesting to note that both VBM and ROI-based studies confirmed that medial temporal lobe is a key area to detect the first signs of AD^20–25^. These studies also showed that HIPP is one of the earliest region altered by AD in the medial temporal lobe^26^. Moreover, HIPP volume is one of the criterion that can be used to confirm the diagnosis of AD in clinical routines^27^. Recently, advanced methods were proposed to capture subtler structural alterations of HIPP^9, 28–30^. Those techniques demonstrated an increase of detection and prediction performances at different AD stages compared to volume-based methods^30^. Among them, patch-based grading (PBG) methods demonstrated competitive results to detect the earliest stages of AD before a clinical diagnosis can be made^9, 29, 31^. The main idea of this approach is to capture inter-subject pattern similarities via non-local comparisons between two groups of subjects. Such methods have shown their ability to predict AD more than seven years before the conversion to dementia^32^ and might help for a differential diagnosis^33^.

Thus, the hippocampus has been one of the most studied structures in order to diagnose AD. However, this structure is complex and not homogeneous. HIPP is subdivided into several subfields, each one having specific characteristics. The terminology differs across segmentation protocol^34^ but the most recognized definition^35^ mainly divides HIPP into the subiculum, the cornu ammonis (CA1/2/3/4), and the dentrate gyrus (DG). The CA1 subfield represents the biggest area in the hippocampus. It is composed by different layers called the stratum radiatum (SR), the stratum lacunosum (SL), the stratum molecular (SM), and the stratum pyramidale (SP). Furthermore, hippocampal subfields are not equally impacted by AD^36–42^. Indeed, several MRI studies demonstrated that subfields are impacted differently according to AD stages. Postmortem, and *in vivo* imaging studies showed that CA1SR-L-M are the subfields impacted with the greatest atrophy in advanced AD^38, 39, 41^. Recently, it has been shown that subiculum is the earliest affected hippocampal region^42, 43^. These studies indicate that a subfield analysis of HIPP alterations at finer scale could provide better tool for AD detection and prediction.

Although structural MRI is a valuable imaging technique to measure global structural modifications, such modality is not able to capture microstructural degradation. However, the microstructural modifications caused by AD are considered to occur before the atrophy measured by structural MRI. Therefore, diffusion MRI (d-MRI) appears as a potential candidate to detect the earliest sign of AD. Several diffusion tensor imaging (DTI) studies proposed automatic methods to detect modifications of diffusion parameters into the whole white matter using machine learning^44–46^. Others studies showed modifications of diffusion parameters for AD patients into specific white matter structures such as corpus callosum^47, 48^, fornix^49^, cingulum^47^ and also in gray matter tissue such as hippocampus^50^. More advanced d-MRI studies using brain connectivity and fiber tracking have been proposed to extract features describing axonal fibers alterations^49, 51, 52^. Finally, it has been shown that hippocampal mean diffusivity (MD) is correlated to pathology progression and thus could be used as an efficient biomarker of AD^53^. Moreover, it was demonstrated that MD increases with the development of AD in the gray matter^54–56^. In a previous work, we showed that patch-based features applied on DTI demonstrated competitive performances to classify the early stages of AD^57^. Methods proposing to fuse d-MRI and s-MRI biomarkers was developed using the complementarity of these two MRI modalities^58, 59^. Recently, a study combining volumetric measurements and mean diffusivity of HIPP subfields demonstrated that CA1 and subiculum are the most impacted in late AD stage^43^. These studies showed the complementarity of s-MRI and d-MRI to capture early alteration led by AD.

All these elements indicate that multimodal analysis of hippocampal subfields using an advanced image analysis framework could provide valuable tool to improve AD detection and prediction. Consequently, in this paper, we propose to study hippocampal subfields efficiencies using s-MRI and d-MRI modalities. We have developed a novel multimodal patch-based grading fusion scheme to better capture such structural and microstructural alterations. First, we compare the performance of our novel method with volume and MD within the whole hippocampus. Second, we demonstrate state-of-the-art performances compared to more advanced d-MRI based methods. Finally, we study the efficiency of hippocampal subfields to improve AD detection and prediction with volume, MD and our multimodal patch-based grading method. Our results demonstrate that the study of hippocampus at finner scale improves AD prediction. Indeed, the experiments show that biomarkers based on whole hippocampus obtain best results for AD detection but biomarkers based on subiculum obtain best results for AD prediction.

## Materials

### Dataset

Data used in this work were obtained from the Alzheimer’s Disease Neuroimaging Initiative (ADNI) dataset (http://adni.loni.ucla.edu). ADNI is a North American campaign launched in 2003 with aims to provide MRI, positron emission tomography scans, clinical neurological measures and other biomarkers. This dataset includes AD patients, MCI and control normal (CN) subjects. The group of MCI is composed of subjects who have abnormal memory dysfunctions. In this work we used data from the ADNI-2 campaign that proposes eMCI and lMCI stages. The eMCI and lMCI subgroups were obtained with the Wechsler Scale-Revised Logistical Memory I and II tests in accordance with the education levels of each subject. ADNI-2 provides T1-weighted (T1w) MRI and DTI scans for 54 CN, 79 eMCI, 39 lMCI and 47 AD subjects. Only patients whose have T1w and DTI were selected in our work. Hence, in this work we used 52 CN, 99 MCI composed of 65 eMCI, 34 lMCI and 38 AD instead of the whole initial ADNI-2 dataset. Table 1 shows the distribution of the data for each group. The s-MRI and d-MRI scans used for all considered subjects in this study were acquired with the same protocol (https://adni.loni.usc.edu/wp-content/uploads/2010/05/ADNI2_GE_3T_22.0_T2.pdf). T1w MRI acquisition protocol had been done with the 3D accelerated sagittal IR-SPGR, according to the ADNI protocol^60^. The d-MRI were composed of 46 separate angles, 5 T2-weighted images with no diffusion sensitization (b0 images) and 41 directions (b=1000s/mm^2^). The d-MRI protocol was chosen to optimize the signal-to-noise ratio in a fixed scan time^61^. The native resolution of s-MRI and d-MRI was set to 1mm^3^ and 2mm^3^, respectively.

**Table 1.** Description of the dataset used in this work. Data are provided by ADNI.

|  | CN | eMCI | IMCI | AD |
| --- | --- | --- | --- | --- |
| Number of subjects | 52 | 65 | 34 | 38 |
| Age (years) | $72.6 \pm 5.9$ | $73.0 \pm 7.7$ | $73.5 \pm 6.6$ | $73.84 \pm 8.7$ |
| Gender (female/male) | 29/23 | 39/26 | 21/13 | 20/18 |
| MMSE | $28.9 \pm 1.2$ | $28.2 \pm 1.5$ | $27.3 \pm 1.8$ | $23.4 \pm 1.7$ |
| CDR-SB | $0.0 \pm 0.1$ | $1.2 \pm 0.6$ | $1.7 \pm 0.8$ | $4.6 \pm 1.4$ |
| RAVLT | $45.4 \pm 9.7$ | $36.5 \pm 10.2$ | $30.7 \pm 8.9$ | $22.6 \pm 7.0$ |
| FAQ | $0.2 \pm 0.9$ | $2.3 \pm 3.7$ | $4.3 \pm 4.8$ | $14.6 \pm 6.6$ |
| ADAS11 | $5.2 \pm 3.0$ | $8.1 \pm 3.6$ | $12.5 \pm 4.9$ | $20.2 \pm 7.6$ |
| ADAS13 | $8.4 \pm 4.4$ | $13.3 \pm 5.4$ | $20.2 \pm 6.7$ | $30.0 \pm 9.0$ |

### MRI processing

T1w images were processed using the volBrain system^62^ (http://volbrain.upv.es). This system is based on an advanced pipeline providing automatic segmentation of different brain structures from T1w MRI. The preprocessing is based on: (a) a denoising step with an adaptive non-local mean filter^63^, (b) an affine registration in the MNI space^64^, (c) a correction of the image inhomogeneities^65^ and (d) an intensity normalization.

Afterwards, segmentation of hippocampal subfields was performed with HIPS^66^ based on a combination of non-linear registration and patch-based label fusion^67^. This method uses a training library based on a dataset composed of high resolution T1w images manually labeled according to the protocol proposed by Winterburn et al. (2013)^35^. To perform the segmentation, the ADNI images are up-sampled with a local adaptive super resolution method to fit in the training image resolution^68^. The method provides automatic segmentation of hippocampal subfields gathered into 5 labels: Subiculum, CA1SP, CA1SR-L-M, CA2-3 and CA4/DG (see Figure 1). Finally, an estimation of the total intra-cranial volume is performed^69^.

**Figure 1.**
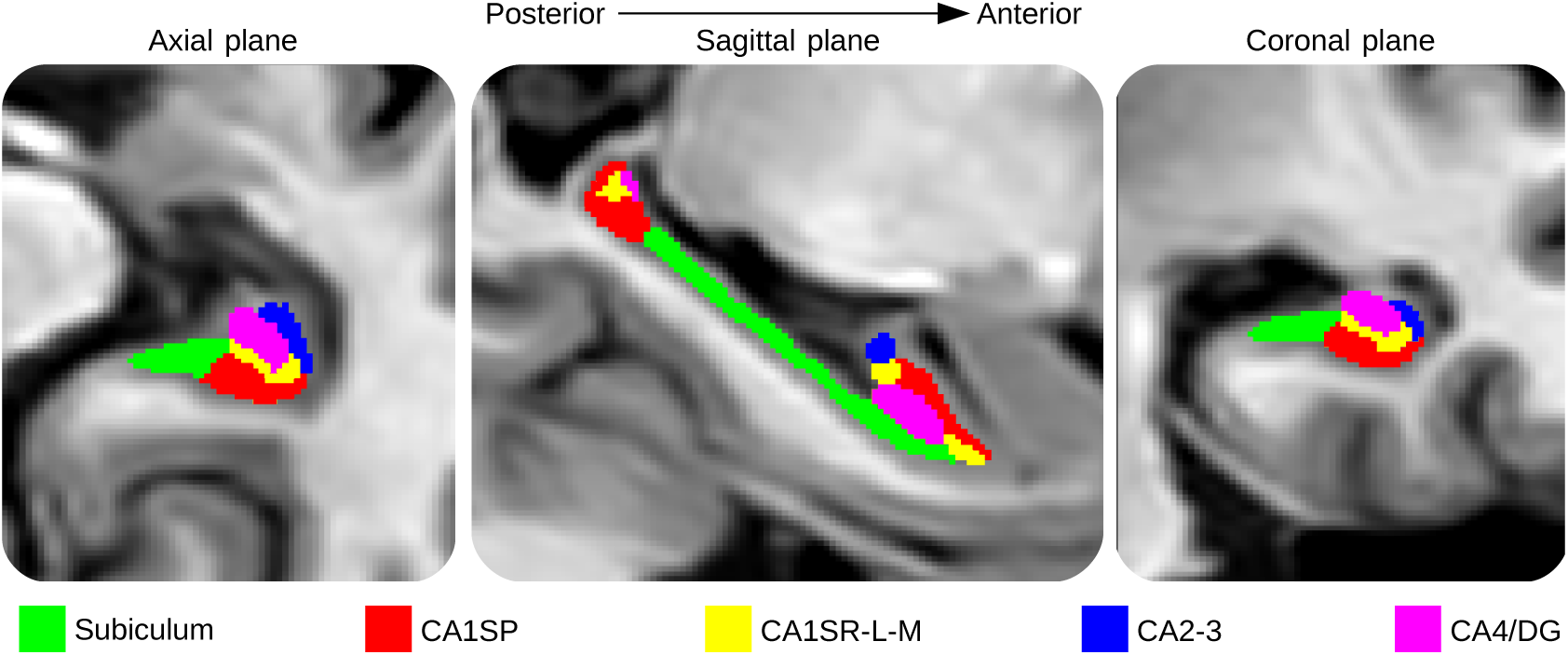
Segmentation of the hippocampal subfields. From left to right, segmentation maps of right hippocampal subfields displayed on the axial, sagittal and coronal plane.

### DTI processing

The preprocessing of the diffusion weighted images is based on: (a) a denoising with a LPCA filter^70^, (b) a correction of the head motion using an affine registration and (c) an affine and a non-rigid registration to the T1w MRI in the MNI space^64^. Afterwards, a diffusion tensor model^71^ is fitted at each voxel using Dipy library^72^. To analyze microstructural modifications, the MD is estimated within each hippocampal subfield and the whole HIPP structure with the segmentation described in the previous section. MD is defined as 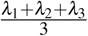 where *λ*_1_, *λ*_2_, *λ*_3_ are the three eigenvalues of the fitted tensor.

Finally, a quality control was proceeded to exclude data presenting miss-segmentation or miss-registration after image preprocessing step.

## Methods

### Patch-based grading

Patch-based grading was firstly proposed for s-MRI^9^. The main idea of this exemplar-based method is to use the capability of patch-based techniques to capture subtle signal modifications related to anatomical degradations caused by AD. To date, PBG methods demonstrate state-of-the-art performances to detect the earliest stage of AD^73^. To determine the pathological status of the subject under study, PBG method estimates at each voxel the state of cerebral tissues by a similarity measurement. This measurement is performed between the anatomical pattern of the subject under study and those extracted from two training populations, one healthy and another one unhealthy.

First, a training library *T* composed of two datasets of images is built: one with images from CN subjects and the other one from AD patients. Next, for each voxel *x*_*i*_ of the region of interest in the considered subject *x*, PBG method produces a weak classifier denoted *g*_*x*_*i*__. This weak classifier provides a surrogate of the pathological grading at the considered position. The weak classifier is computed using a measurement of the similarity between the patch *P*_*x*_*i*__ surrounding the voxel *x*_*i*_ belonging to the image under study and a set *K*_*x*_*i*__ of the closest patches extracted from the library *T*. The most similar patches are found using an approximative nearest neighbor method^74^. The grading value *g*_*x*_*i*__ at *x*_*i*_ is defined as:

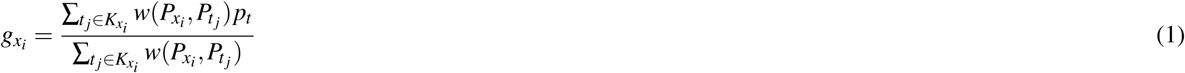

where *P*_*t*_*j*__ is the patch surrounding the voxel *j* belonging to the training template *t ∈ T*. *w*(*x*_*i*_, *t*_*j*_) is the weight assigned to the pathological status *p*_*t*_ of the training image *t*. We estimate *w* such as:

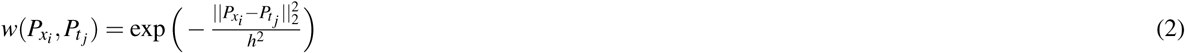

where 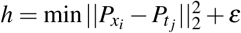 and *ε* → 0. The pathological status *p*_*t*_ is set to −1 for patches extracted from AD patient and to 1 for patches extracted from CN subject. Therefore, PBG method provides at each voxel a score representing an estimation of the alterations caused by AD. Consequently, cerebral tissues strongly altered by AD have grading values close to −1 contrary to healthy one with scores close to 1.

### Multimodal patch-based grading fusion

Patch-based method presented in the previous section was firstly designed to capture structural alterations in T1w MRI. Recently, we proposed to extent this method to DTI modality in order to detect microstructural modifications^57^. We showed the efficiency of MD grading to improve the classification of the early stages of AD.

In this study, we propose a new framework to perform multimodal patch-based grading (MPBG). To this end, we developed adaptive fusion of grading maps derived from different modalities (see example of grading maps on Fig. 2). As shown in the following, this fusion provides more robust and accurate biomarkers compared to monomodal PBG biomarkers.

**Figure 2.**
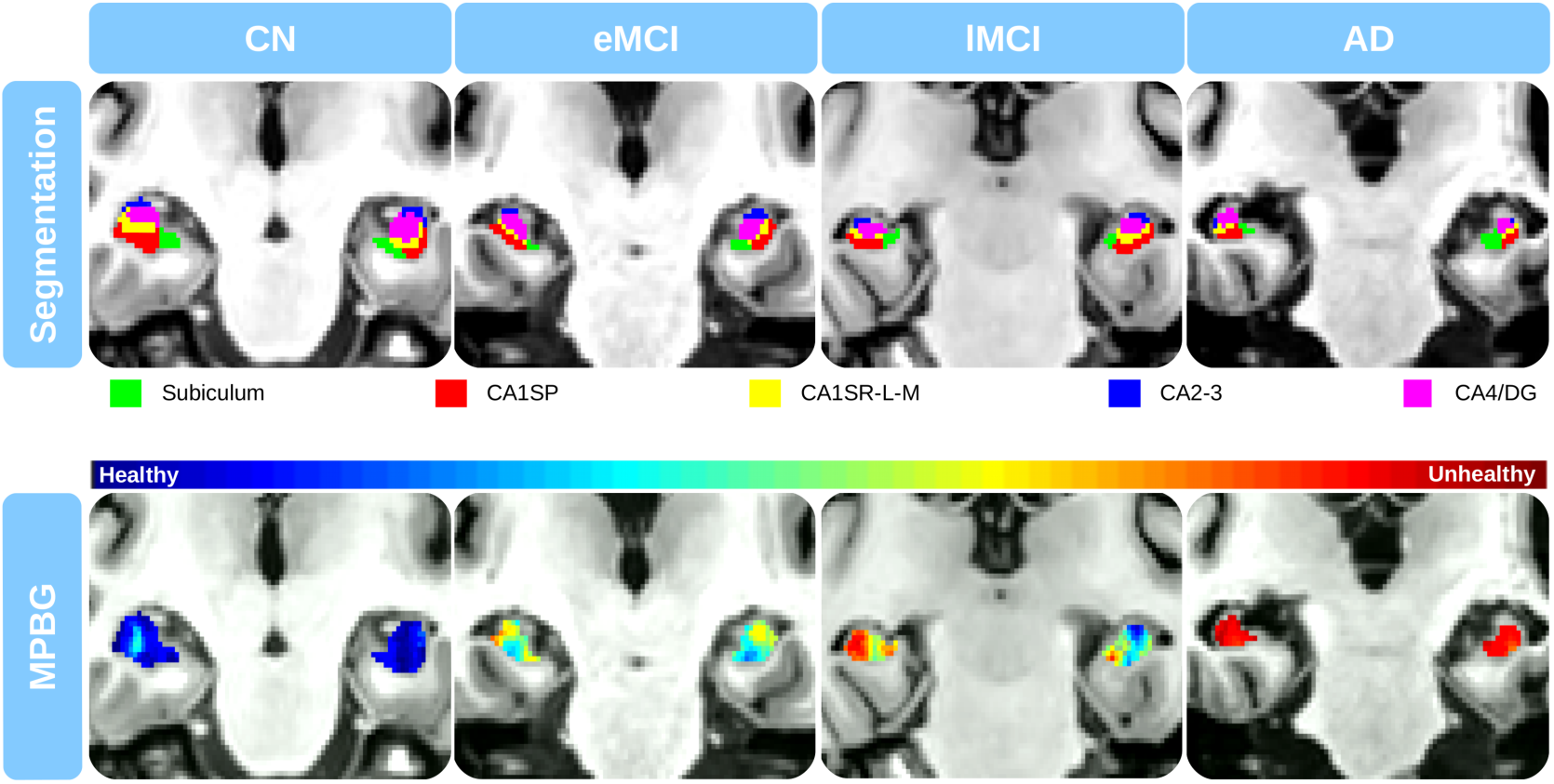
Results obtained for different severities of cognitive impairments. From top to bottom slices on the coronal plane of the segmentation maps and the fusion of T1w and MD patch-based grading with the proposed multimodal patch-based grading method. The blue and the red colors represent the healthy and altered tissues, respectively.

First, as in previous section, for each modality a training library of CN and AD subjects is built. Next, at each voxel within the ROI of the considered subject and for each modality, a set *K* of most similar patches is extracted. This step provides one set *K* of patches per modality *m ∈ M*, where *M* corresponds to the set of the different modalities provided. Nevertheless, at each voxel the quality of the grading estimation is not the same for all the modalities. Therefore, the degree of confidence is estimated with the function *α* defined as:

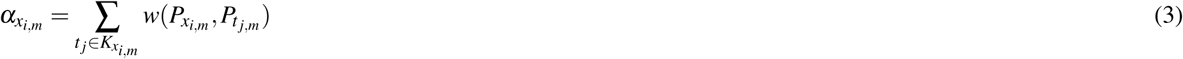

that reflects the confidence of the grading value *g*_*x*_*i*__ for the modality *m* at the voxel *x*_*i*_. This confidence measure is derived from multi-feature fusion^75^. Thus, each modality provides a weak classifier at each voxel that is weighted with its degree of confidence *α*_(_*x*_*i,m*_). The multimodal grading, denoted *g*_*x*_*i*__, is given by:

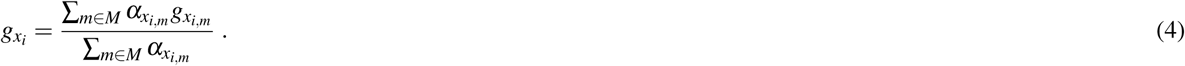

In other words, the weight *w* and *K*_*x*_*i,m*__ are estimated independently for each modality and combined afterwards. Therefore, the proposed combination framework is spatially adaptive and takes advantage of having access to a local degree of confidence *α*_*x*_*i,m*__ for each modality *m*. Basically, when the set of patches found for a modality in the training library is composed of good candidates (*i.e.*, patches very similar to the patch from the subject under study), our confidence *α*_*x*_*i,m*__ in the grading estimation for this modality is high. At the end, this modality will have a high weight in the mixing procedure described in (4).

### Features estimation

Features were estimated in each hippocampal subfield and over the whole hippocampus as the union of all hippocampal subfields masks. To reduce the inter-individual variability, all volumes are normalized by the total intra-cranial volume^76^. Afterwards, we aggregate local weak classifiers of the grading map into a single feature for each considered structure (*i.e.*, hippocampal subfields and whole HIPP) by averaging them. Therefore, patch-based grading features are computed by an unweighted vote of the weak classifiers using the segmentation masks (see Fig. 3). Finally, to prevent the bias introduced as the structure alterations due to aging, all the features (*i.e.*, volume, mean of MD and MPBG) are age corrected with a linear regression based on the CN group^77^.

**Figure 3.**
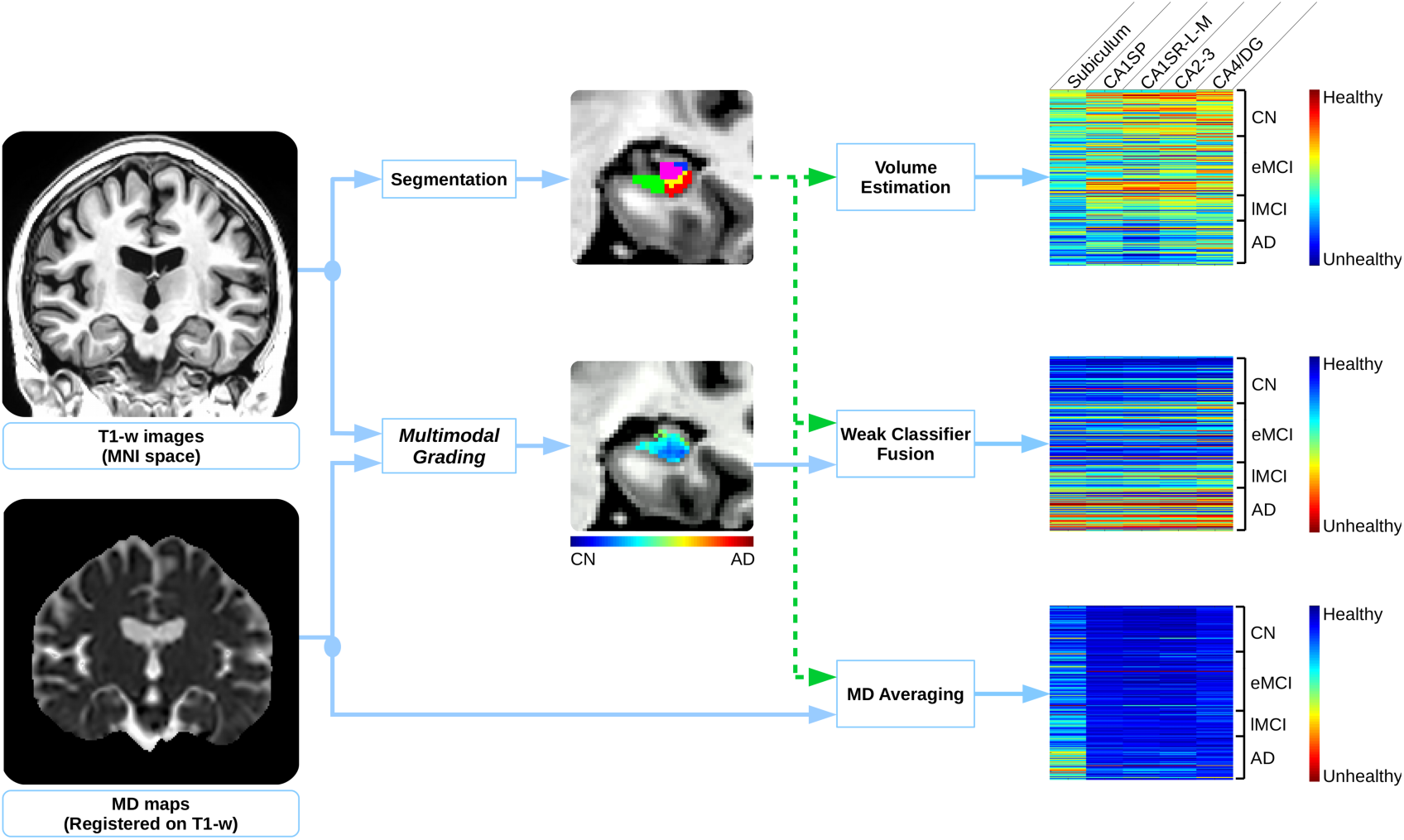
Proposed multimodal patch-based grading framework. At left, the input data: T1w images registered into the MNI space and MD maps registered on the T1w images. At the middle: a coronal view of hippocampal subfields segmentation on T1w, and the corresponding coronal view of a MPBG map estimated on T1w and MD. At right, the considered subfield biomarkers for all subjects under study. From top to bottom, the features are the volumes, the MPBG values, and the average of MD.

### Implementation

To find the most similar patches in the training library, we use the OPAL method^78^. OPAL is a fast approximate nearest neighbor patch search technique. This method enables to process each modality in about 4 seconds on a standard computer. The training library is equally composed of 37 images for both CN and AD subjects, leading to |*T*| = 76. The number of patches extracted from each the training library is *K* = 160 (*i.e.*, 80 from CN subjects and 80 from AD patients) and the patch size is 5 × 5 × 5 voxels. Furthermore, as done in our PBG DTI study^57^, we used zero normalized sum of squared differences for T1w to compute the L2 norm (see Equation (2)). On the other hand, d-MRI is a quantitative imaging technique. Therefore, to preserve the quantitative information, a straight sum of squared differences is used for MD in Equation (2),

### Validation

To evaluate the efficiency of each considered biomarker to detect AD alterations, CN group is compared to AD patients group. In addition, to discriminate the impairment severity of MCI group, eMCI versus lMCI classification is conducted. The classification step is performed with a linear discriminant analysis (LDA) within a repeated stratified 5-fold cross-validation iterated 200 times. Mean area under the curve (AUC) and mean accuracy (ACC) are computed to compare performance for each biomarker over the 200 iterations.

## Results

In this section, the results are presented in three parts. In the first part, we compare the different approach applied within the entire HIPP structure to evaluate the performance of our new MPBG compared to usual biomarkers such as volume and average MD. Afterwards, in the second part, we compare the accuracy of each considered biomarkers within hippocampal subfields to investigate the potential of hippocampal subfields analysis to improve result of AD detection and prediction. Finally, in a last part, we compare the results of our proposed multimodal biomarker with state-of-the-art methods based on d-MRI to show the competitive performance of our approach.

### Whole hippocampus

Results of the comparisons over the whole HIPP are represented in Table 2. In this experiment, we compared the results of volume, mean of MD and PBG applied with both modality and MPBG over the whole hippocampus.

**Table 2.** Mean AUC of the different features estimated over the whole hippocampal structure. In bold font the best result for each specific comparison. All results are expressed in percent.

| Method | CN vs. AD | eMCI vs. IMCI |
| --- | --- | --- |
| Volume | 86.6 | 59.4 |
| MD | 80.6 | 55.6 |
| T1w PBG | **92.6** | 67.5 |
| MD PBG | 89.2 | **69.5** |
| MPBG | 92.1 | **69.5** |

First, the hippocampus volume and its average of MD were compared. For CN versus AD classification the volume obtains 86.6% of AUC and the average of MD obtains 80.6%. For eMCI versus lMCI classification the volume and the average of MD obtain 59.4% and 55.6% of AUC, respectively. Experiments demonstrate that the hippocampus the volume obtains better classification results than the average of MD for all comparison, especially for CN versus AD. Second, PBG biomarkers applied with T1w and MD were compared. The results showed that T1w PBG provides better results than MD PBG with 92.6% of AUC for CN versus AD classification. However, for eMCI versus lMCI classification MD grading provides the best results with 69.5% of AUC. MPBG methods combining both modalities reaches the best results for CN versus AD and eMCI versus lMCI with 92.1% and 69.5% of AUC, respectively. Finally, the proposed MPBG biomarker provides results similar to the best modalities for all considered comparisons. Compared to volume, MPBG improves CN versus AD comparison result by 5.5% of AUC and by over 10% of AUC for eMCI versus lMCI comparison. Thus, MBPG biomarker has a good capability to capture modifications caused by AD at different severity stages (see Figure 2).

### Hippocampal subfields

Figure 4 shows the distribution of volumes (A), average of MD (B) and MPBG (C) for each hippocampal subfield at each different AD stages. For each comparison a p-value was estimated with a multi-comparison test^79^. We can note that for all hippocampal subfields, alterations caused by the disease are related to a volume and MPBG decrease with a MD increase. Subiculum subfield presents the most significant differences for several comparisons. More importantly, it is the only subfield providing a p-value inferior than 0.05 for the comparison CN versus eMCI with volume, a p-value inferior than 0.01 to lMCI versus AD with MD and a p-value inferior to 0.001 to eMCI versus lMCI with MPBG, which are the most challenging comparisons. The distribution of MPBG shows a better discrimination between each group for all hippocampal subfields. Indeed, MPBG applied within CA1SP, and CA1SR-L-M provides p-values inferior than 0.01 for eMCI versus lMCI. Moreover, MPBG applied within the subiculum provides p-value inferior than 0.001 for the same comparison. Thus, MPBG enables to perform a detection of AD with each subfield with an advantage for subiculum for the comparison of eMCI versus lMCI.

**Figure 4.**
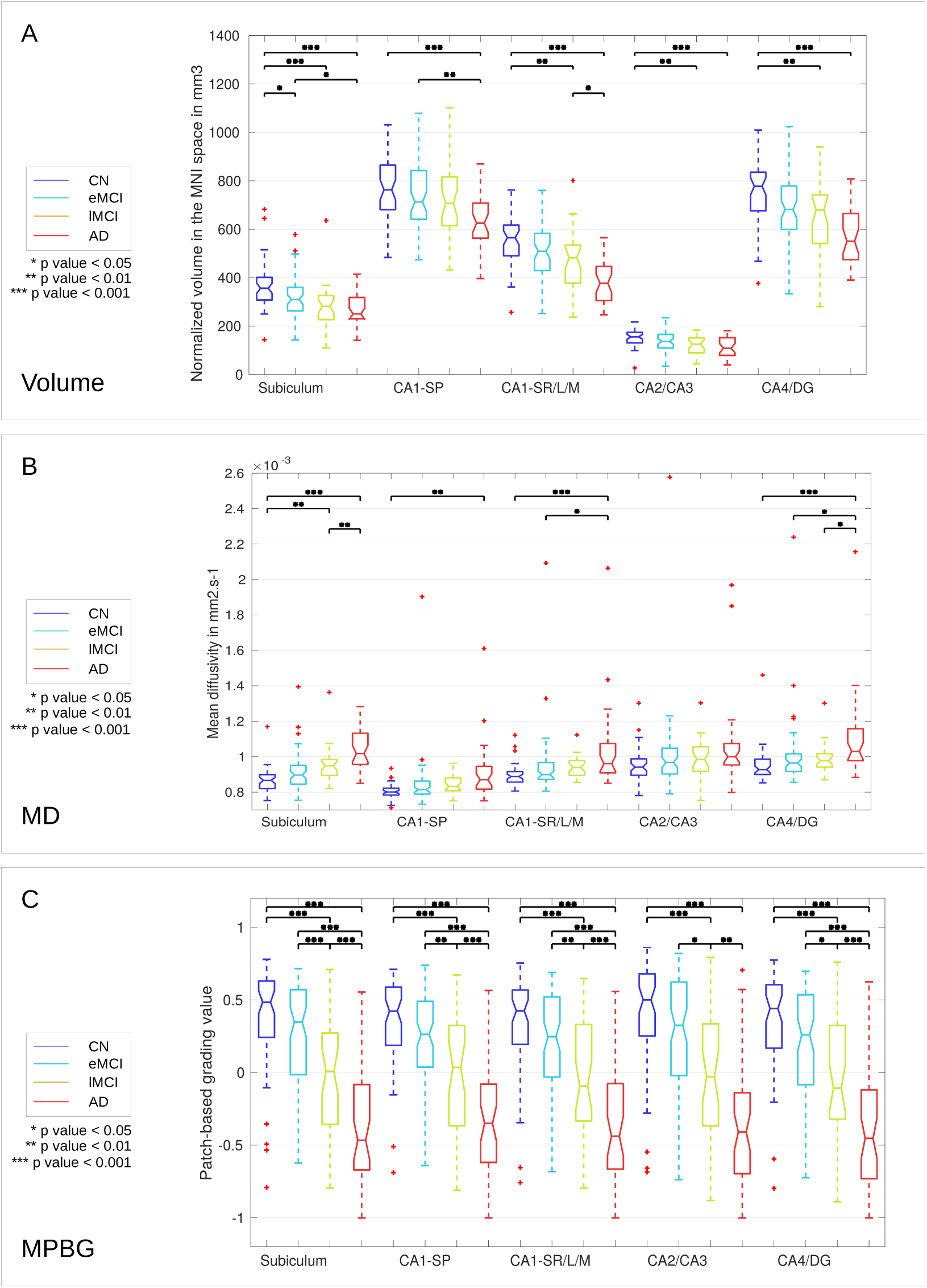
Distribution of the volume (A), MD (B) and MPBG (C) for the different considered groups. The normalized volumes are provided in *mm*^3^ in the MNI space for each subfield, MD is the mean of MD values into each subfield in *mm*^2^*.s*^−1^, and MPBG is the mean patch-based grading values into each subfield. Blue, cyan, orange and red colors represent CN, eMCI, lMCI and AD subjects, respectively. Statistical tests have been performed with ANOVA procedure and corrected for multiple comparisons with the Bonferroni’s method. The p-values inferior to 0.05, inferior to 0.01, and inferior to 0.001 are represented with *, **, and ***, respectively.

To estimate the efficiency of the considered biomarkers for AD detection, we also performed a classification experiment. Figures 5 shows the results of two comparisons, CN versus AD (part noted A in the figure) and eMCI versus lMCI (part noted B). First, for AD diagnosis (*i.e.*, CN versus AD classification), the subfield providing the most discriminant volume is the CA1S-R-L-M with an AUC of 86.0%. Moreover, the most discriminant MD biomarker is given by the subiculum with an AUC of 88.1%. For this comparison, MD of subiculum is the only biomarker performing better results than whole hippocampus. The best results obtained by MPBG feature is provided by the CA1SP with an AUC of 92.1% followed by CA1S-R-L-M and subiculum. Second, for eMCI versus lMCI classification, the subiculum provides the best results for each considered feature. Indeed, subiculum obtained an AUC of 66.1% for the volume, 62.4% for the average of MD, and 71.8% for MPBG. Moreover, subiculum provided better results than whole hippocampus for each feature. Thus, the experiments conducted with three different biomarkers showed that the use of hippocampal subfields, especially the subiculum, enables to obtain better results for AD prediction than the whole hippocampal analysis.

**Figure 5.**
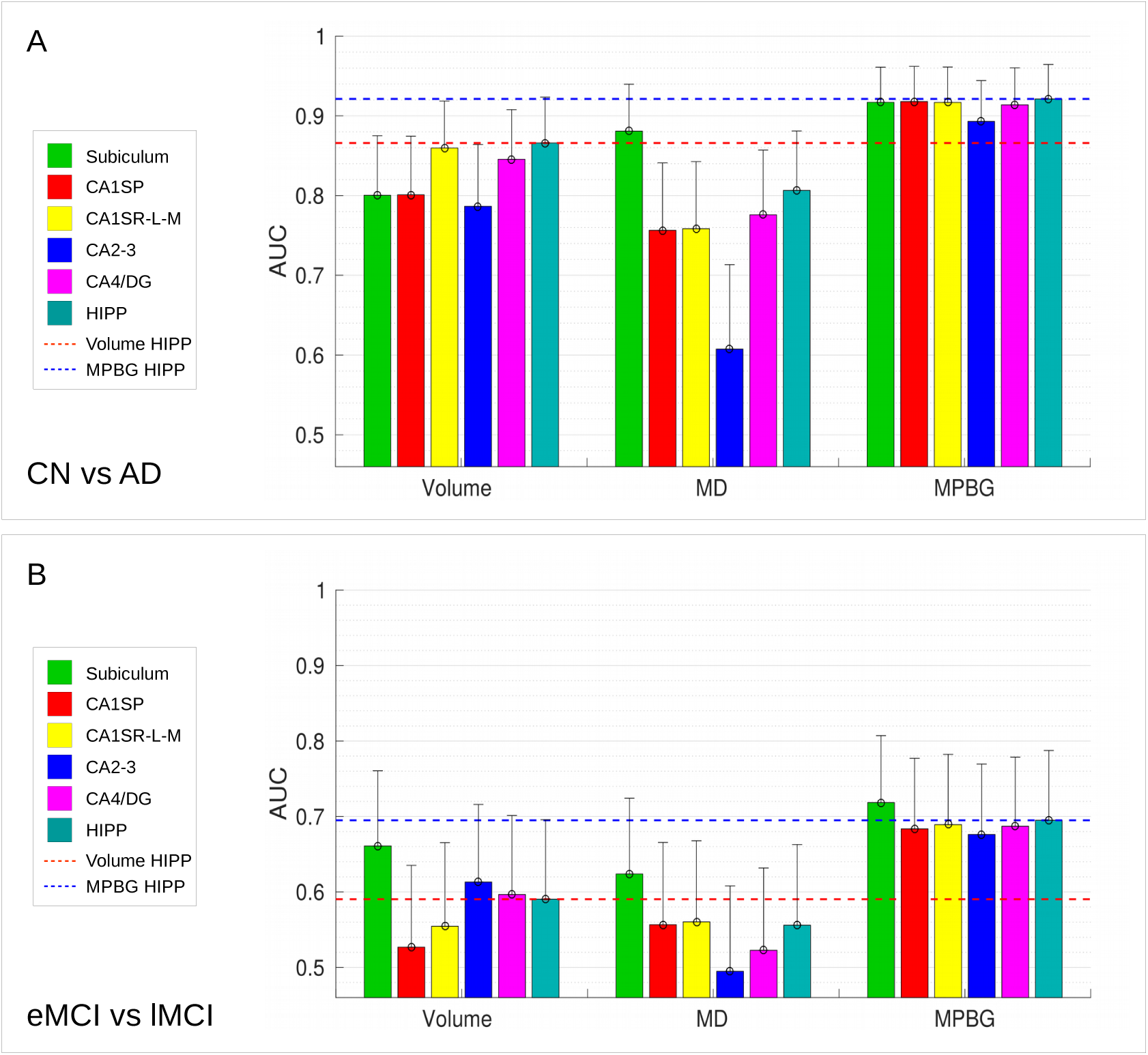
AUC computed for CN versus AD (A), eMCI versus lMCI (B) comparisons with the different considered biomarkers in each hippocampal area. Results of subfields are grouped by features (*i.e.*, the volume, the average of MD and the MPBG). Upper bounds of confidence interval are represented with vertical bars. Whole HIPP volume biomarker provides the best results with a mean AUC of 86.6% for CN versus AD comparison, followed by the CA1S-R-L-M volume that obtains a mean AUC of 86%. Subiculum volume provides the best results for eMCI versus lMCI with a mean AUC of 66.1%. The average of MD for subiculum obtains the best results for CN versus AD and eMCI versus lMCI with a mean AUC of 88.0% and 62.4%, respectively. Whole HIPP MPBG obtains the best results for CN versus AD with a mean AUC of 92.1%. Subiculum MPBG obtains the best results for eMCI versus lMCI comparison with a mean AUC of 71.8%. This comparison shows that subiculum is the only biomarker providing better results than whole HIPP.

### Comparison with state-of-the-art methods

To evaluate the performance of the proposed MPBG, we compared it with state-of-the-art multimodal methods using d-MRI. To this end, we used the ACC values published by the authors. Table 3 shows the comparison of our proposed biomarkers within the hippocampal area providing the best results (*i.e.* the whole HIPP and the subiculum) with the state-of-the-arts methods using similar dataset based on ADNI-2. We compared these biomarkers with a method using features based on tractography^80^, a method based on a connectivity network of the different brain structures^52^, and a voxel-based method that analyzes alterations of white matter^81^. The results of comparison show that MPBG over whole HIPP obtains the best score for AD versus CN with 88.1% of accuracy while the best result is achieved by a voxel-based method with a feature selection^81^ that obtained 87.0% on similar ADNI2 dataset. For the best of our knowledge, the only work providing eMCI and lMCI comparison^52^ using d-MRI from similar ADNI2 dataset is based on a connectivity network and obtained 63.4%. These comparisons demonstrate the relevance of MPBG biomarkers for AD detection and prediction. Indeed, our method provides similar results than the best methods with similar dataset for CN versus AD classification and provides the best results for eMCI versus lMCI classification.

**Table 3.** Comparison of our proposed MPBG biomarkers with state-of-the-arts methods based on d-MRI using a similar ADNI2 dataset. All results are expressed in percentage of accuracy.

| Method | Feature | Classifier | CN | eMCI | IMCI | AD | CN vs. AD | eMCI vs. IMCI |
| --- | --- | --- | --- | --- | --- | --- | --- | --- |
| Nir et al. (2015)<sup>80</sup> | Tractography | SVM | 44 | 74 | 39 | 23 | 84.9% | n/a |
| Prasad et al. (2015)<sup>52</sup> | Connectivity network | SVM | 50 | 74 | 38 | 38 | 78.2% | 63.4% |
| Maggipinto et al. (2017)<sup>81</sup> | Voxel-based | RF | 50 | 22 | 18 | 50 | 87.0% | n/a |
| MPBG HIPP | Patch-based | LDA | 62 | 65 | 34 | 38 | **88.1%** | 68.8% |
| MPBG Subiculum | Patch-based | LDA | 62 | 65 | 34 | 38 | 86.5% | **70.8%** |

Moreover, the proposed MPBG method based on subiculum improves the performance for eMCI versus lMCI classification with an accuracy of 70.8%, that increases by 2% the accuracy based the whole HIPP and over 6% compared to a connectivity network based method.

## Discussion

In this work, a multimodal analysis of the hippocampal subfields alterations caused by AD is proposed. First, the structural and micro-structural alterations were captured from two MRI modalities with different methods. Thus, the use of volume, MD, and the proposed MPBG methods were investigated to achieve this analysis. In this section, the efficiency of these different methods applied into the whole hippocampus and each hippocampal subfield are discussed.

### Whole hippocampus biomarkers

We first compared the performance of different methods applied to the whole hippocampus (see Table 2). The experiments showed that volume and mean of MD within a structure as the hippocampus does not provides discriminant biomarkers to detect early stages of AD. The MPBG method based on s-MRI and d-MRI obtains best results compared to the volume and the average of MD. Moreover, compared to recent methods proposed for AD detection^82^ (see Table 3), proposed MPBG demonstrates state-of-the-art performances for AD detection and prediction. These results emphasize the relevance of using more accurate biomarker, such MPBG to study the effectiveness of hippocampal subfields for AD detection and prediction.

### Hippocampal subfield biomarkers

The main contribution of this study is the multimodal analysis of hippocampal subfields. Indeed, most of the proposed biomarkers based on hippocampus focused only on the whole structure or study alterations of hippocampal subfields with methods that do not provide sensitive biomarkers to detect early modification caused by AD. The lack of work studying alterations of hippocampal subfields with advanced biomarkers could be explained by the fact that automatic segmentation of the hippocampal subfields is a complex task due to subtle borders dividing each area.

In this work, we compared the efficiency of diffusion MRI and multimodal patch-based biomarkers for AD detection and prediction over the hippocampal subfields. Comparisons based on MD, volume and multimodal patch-based biomarkers showed that the subiculum is the most discriminant structure in the earliest stage of AD providing the best results for AD prediction (see Figure 4 and 5). However, whole hippocampus structure, followed by CA1SR-L-M, obtains best results for AD detection.

These results are in accordance with literature studies based on animal model and *in vivo* imaging combining volume and MD demonstrating that subiculum is the earliest hippocampal region affected by AD^42, 43^. Moreover, postmortem studies showed that the hippocampal degeneration in early stages of AD is not uniform. After the apparition of alterations in the EC, the pathology spreads to the subiculum, CA1, CA2-3 and finally the CA4 and DG subfields^36, 37, 42, 83^. It is interesting to note that the results of our experiments using volume-based biomarkers are also coherent with the previous *in-vivo* imaging studies that analyzed the atrophy of each hippocampal subfield at advanced stage of AD. These studies showed that CA1 is the subfield impacted by the strongest atrophy^38, 39, 84, 85^. Furthermore, studies using ultra-high field at 7T enabling CA1 layers discrimination showed that CA1SR-L-M is the subfields showing the greater atrophy at advanced stages of AD^40, 41^.

## Conclusion

In this paper, we analyzed hippocampal subfield alterations with a multimodal framework based on structural and diffusion MRI. In addition, to study tenuous modifications occurring into each hippocampal subfield, we developed a new multimodal patch-based framework using T1w and DTI. Our novel MPBG method were compared to the volume and the average of MD over the whole hippocampus. This comparison demonstrated that MPBG method improves performances for AD detection and prediction. In addition, a comparison with state-of-the-art diffusion-based methods showed the competitive performance of MPBG biomarkers. Finally, an analysis of the hippocampal subfields with the volume, the average of MD and MBPG methods was conducted. Although CA1 is the subfields having the greater atrophy in the late stage of AD, the experiments demonstrated that whole hippocampus provides best biomarker for AD detection while subiculum provides best biomarker for AD prediction.

## Acknowledgements

This study has been carried out with financial support from the French State, managed by the French National Research Agency (ANR) in the frame of the Investments for the future Program IdEx Bordeaux (HL-MRI ANR-10-IDEX-03-02), Cluster of excellence CPU and TRAIL (HR-DTI ANR-10-LABX-57) and the CNRS multidisciplinary project “Défi imag’In”.

Data collection and sharing for this project was funded by the Alzheimer’s Disease Neuroimaging Initiative (ADNI) (National Institutes of Health Grant U01 AG024904) and by the National Institute on Aging, the National Institute of Biomedical Imaging and Bioengineering, and through generous contributions from the following: AbbVie, Alzheimer’s Biogen; Bristol-Myes Squibb Company; CereSpir, Inc.; Cogstate; Eisai Inc.; Elan Pharmaceuticals, Inc.; Eli Lilly and Company; EuroImmun; F. Hoffman-La Roche Ltd and its affiliated company Genentech, Inc.; Fujirebio; GE Healthcare; IXICO Ltd.; Janssen Pharmaceutical Research & Development LLC.; NeuroRx Research; Neurotrack Technologies; Novartis Pharmaceuticals Corporation; Pfizer Inc.; Piramal Imaging; Servier; Takeda Pharmaceutical providing funds to support ADNI clinical sites in Canada. Private sector contributions are facilitated by the Foundation for the National Institutes of Health (www.fnih.org). The grantee organization is the Northern California Institute of Research and Education, and the study is coordinated by the Alzheimer’s Therapeutic Research Institute at the University of Southern California. ADNI data are disseminated by the Laboratory for Neuro Imaging at the University of Southern California.

## References

1. Petersen, R. C. et al. Current concepts in mild cognitive impairment. Arch. neurology 58, 1985–1992 (2001).

2. Aisen, P. S. et al. Clinical core of the Alzheimer’s Disease Neuroimaging Initiative: progress and plans. Alzheimer’s Dementia 6, 239–246 (2010).

3. Bron, E. E. et al. Standardized evaluation of algorithms for computer-aided diagnosis of dementia based on structural MRI: The CADDementia challenge. NeuroImage 111, 562–579 (2015).

4. Hyman, B. T., Van Hoesen, G. W., Damasio, A. R. & Barnes, C. L. Alzheimer’s disease: cell-specific pathology isolates the hippocampal formation. Sci. 225, 1168–1170 (1984).

5. West, M. J., Coleman, P. D., Flood, D. G., Troncoso, J. C. et al. Differences in the pattern of hippocampal neuronal loss in normal ageing and Alzheimer’s disease. The Lancet 344, 769–772 (1994).

6. Braak, H. & Braak, E. Staging of Alzheimer’s disease-related neurofibrillary changes. Neurobiol. aging 16, 271–278 (1995).

7. Gómez-Isla, T. et al. Profound loss of layer ii entorhinal cortex neurons occurs in very mild Alzheimer’s disease. J. Neurosci. 16, 4491–4500 (1996).

8. Du, A. et al. Magnetic resonance imaging of the entorhinal cortex and hippocampus in mild cognitive impairment and Alzheimer’s disease. J. Neurol. Neurosurg. & Psychiatry. 71, 441–447 (2001).

9. Coupé, P. et al. Scoring by nonlocal image patch estimator for early detection of Alzheimer’s disease. NeuroImage: clinical 1, 141–152 (2012).

10. Jack, C. R. et al. Medial temporal atrophy on MRI in normal aging and very mild Alzheimer’s disease. Neurol. 49, 786–794 (1997).

11. Ross, S. et al. Progressive biparietal atrophy: an atypical presentation of Alzheimer’s disease. J. Neurol. Neurosurg. & Psychiatry. 61, 388–395 (1996).

12. Kaida, K.-i., Takeda, K., Nagata, N. & Kamakura, K. Alzheimer’s disease with asymmetricx parietal lobe atrophy: a case report. J. neurological sciences 160, 96–99 (1998).

13. Jack, C. R., Petersen, R. C., O’brien, P. C. & Tangalos, E. G. Mr-based hippocampal volumetry in the diagnosis of Alzheimer’s disease. Neurol. 42, 183–183 (1992).

14. Jack, C. R. et al. Hypothetical model of dynamic biomarkers of the Alzheimer’s pathological cascade. The Lancet Neurol. 9, 119–128 (2010).

15. Scher, A. et al. Hippocampal shape analysis in Alzheimer’s disease: a population-based study. Neuroimage 36, 8–18 (2007).

16. Achterberg, H. C. et al. Hippocampal shape is predictive for the development of dementia in a normal, elderly population. Hum. brain mapping 35, 2359–2371 (2014).

17. Fischl, B. & Dale, A. M. Measuring the thickness of the human cerebral cortex from magnetic resonance images. Proc. Natl. Acad. Sci. 97, 11050–11055 (2000).

18. Eskildsen, S. F. et al. Prediction of Alzheimer’s disease in subjects with mild cognitive impairment from the ADNI cohort using patterns of cortical thinning. Neuroimage 65, 511–521 (2013).

19. Ashburner, J. & Friston, K. J. Voxel-based morphometry—the methods. Neuroimage 11, 805–821 (2000).

20. Good, C. D. et al. Automatic differentiation of anatomical patterns in the human brain: validation with studies of degenerative dementias. Neuroimage 17, 29–46 (2002).

21. Karas, G. et al. Global and local gray matter loss in mild cognitive impairment and Alzheimer’s disease. Neuroimage 23, 708–716 (2004).

22. Hirata, Y. et al. Voxel-based morphometry to discriminate early Alzheimer’s disease from controls. Neurosci. letters 382, 269–274 (2005).

23. Klöppel, S. et al. Automatic classification of MR scans in Alzheimer’s disease. Brain 131, 681–689 (2008).

24. Ferreira, L. K., Diniz, B. S., Forlenza, O. V., Busatto, G. F. & Zanetti, M. V. Neurostructural predictors of Alzheimer’s disease: a meta-analysis of VBM studies. Neurobiol. aging 32, 1733–1741 (2011).

25. Wolz, R. et al. Multi-method analysis of MRI images in early diagnostics of Alzheimer’s disease. PloS one 6, e25446 (2011).

26. Frisoni, G. B., Fox, N. C., Jack, C. R., Scheltens, P. & Thompson, P. M. The clinical use of structural MRI in Alzheimer disease. Nat. Rev. Neurol. 6, 67–77 (2010).

27. Hill, D. L. et al. Coalition against major diseases/european medicines agency biomarker qualification of hippocampal volume for enrichment of clinical trials in predementia stages of Alzheimer’s disease. Alzheimer’s & Dementia 10, 421–429 (2014).

28. Gerardin, E. et al. Multidimensional classification of hippocampal shape features discriminates Alzheimer’s disease and mild cognitive impairment from normal aging. Neuroimage 47, 1476–1486 (2009).

29. Tong, T. et al. Multiple instance learning for classification of dementia in brain MRI. Med. image analysis 18, 808–818 (2014).

30. Sørensen, L. et al. Differential diagnosis of mild cognitive impairment and Alzheimer’s disease using structural MRI cortical thickness, hippocampal shape, hippocampal texture, and volumetry. NeuroImage: Clin. (2016).

31. Liu, M., Zhang, D., Shen, D. & Alzheimer’s Disease Neuroimaging Initiative. Ensemble sparse classification of Alzheimer’s disease. NeuroImage 60, 1106–1116 (2012).

32. Coupé, P. et al. Detection of Alzheimer’s disease signature in MR images seven years before conversion to dementia: Toward an early individual prognosis. Hum. brain mapping 36, 4758–4770 (2015).

33. Koikkalainen, J. et al. Differential diagnosis of neurodegenerative diseases using structural MRI data. NeuroImage: Clin. 11, 435–449 (2016).

34. Yushkevich, P. A. et al. Quantitative comparison of 21 protocols for labeling hippocampal subfields and parahippocampal subregions in in vivo MRI: towards a harmonized segmentation protocol. Neuroimage 111, 526–541 (2015).

35. Winterburn, J. L. et al. A novel in vivo atlas of human hippocampal subfields using high-resolution 3T magnetic resonance imaging. Neuroimage 74, 254–265 (2013).

36. Braak, E. & Braak, H. Alzheimer’s disease: transiently developing dendritic changes in pyramidal cells of sector CA1 of the ammon’s horn. Acta neuropathologica 93, 323–325 (1997).

37. Braak, H., Alafuzoff, I., Arzberger, T., Kretzschmar, H. & Del Tredici, K. Staging of Alzheimer disease-associated neurofibrillary pathology using paraffin sections and immunocytochemistry. Acta neuropathologica 112, 389–404 (2006).

38. Apostolova, L. G. et al. Conversion of mild cognitive impairment to Alzheimer disease predicted by hippocampal atrophy maps. Arch. neurology 63, 693–699 (2006).

39. La Joie, R. et al. Hippocampal subfield volumetry in mild cognitive impairment, Alzheimer’s disease and semantic dementia. NeuroImage: Clin. 3, 155–162 (2013).

40. Kerchner, G. et al. Hippocampal CA1 apical neuropil atrophy in mild Alzheimer disease visualized with 7-T MRI. Neurol. 75, 1381–1387 (2010).

41. Kerchner, G. A. et al. Hippocampal CA1 apical neuropil atrophy and memory performance in Alzheimer’s disease. Neuroimage 63, 194–202 (2012).

42. Trujillo-Estrada, L. et al. Early neuronal loss and axonal/presynaptic damage is associated with accelerated amyloid-*β* accumulation in a*β* pp/ps1 Alzheimer’s disease mice subiculum. J. Alzheimer’s Dis. 42, 521–541 (2014).

43. Li, Y.-D., Dong, H.-B., Xie, G.-M. & Zhang, L.-j. Discriminative analysis of mild Alzheimer’s disease and normal aging using volume of hippocampal subfields and hippocampal mean diffusivity: an in vivo magnetic resonance imaging study. Am. J. Alzheimer’s Dis. & Other Dementias 28, 627–633 (2013).

44. O’Dwyer, L. et al. Using support vector machines with multiple indices of diffusion for automated classification of mild cognitive impairment. PloS one 7, e32441 (2012).

45. Dyrba, M. et al. Robust automated detection of microstructural white matter degeneration in Alzheimer’s disease using machine learning classification of multicenter DTI data. PloS one 8, e64925 (2013).

46. Dyrba, M. et al. Predicting prodromal Alzheimer’s disease in subjects with mild cognitive impairment using machine learning classification of multimodal multicenter diffusion-tensor and magnetic resonance imaging data. J. Neuroimaging 25, 738–747 (2015).

47. Nir, T. M. et al. Effectiveness of regional DTI measures in distinguishing Alzheimer’s disease, MCI, and normal aging. NeuroImage: clinical 3, 180–195 (2013).

48. Wang, Z. et al. Interhemispheric functional and structural disconnection in Alzheimer’s disease: a combined resting-state fMRI and DTI study. PLoS One 10, e0126310 (2015).

49. Liu, Y. et al. Diffusion tensor imaging and tract-based spatial statistics in Alzheimer’s disease and mild cognitive impairment. Neurobiol. aging 32, 1558–1571 (2011).

50. Rose, S. E., Andrew, L. & Chalk, J. B. Gray and white matter changes in Alzheimer’s disease: a diffusion tensor imaging study. J. Magn. Reson. Imagin. 27, 20–26 (2008).

51. Wee, C.-Y. et al. Identification of MCI individuals using structural and functional connectivity networks. Neuroimage 59, 2045–2056 (2012).

52. Prasad, G. et al. Brain connectivity and novel network measures for Alzheimer’s disease classification. Neurobiol. aging 36, S121–S131 (2015).

53. Fellgiebel, A. & Yakushev, I. Diffusion tensor imaging of the hippocampus in MCI and early Alzheimer’s disease. J. Alzheimer’s Dis. 26, 257–262 (2011).

54. Kantarci, K. et al. DWI predicts future progression to Alzheimer disease in amnestic mild cognitive impairment. Neurol. 64, 902–904 (2005).

55. Müller, M. J. et al. Functional implications of hippocampal volume and diffusivity in mild cognitive impairment. Neuroimage 28, 1033–1042 (2005).

56. Fellgiebel, A. et al. Predicting conversion to dementia in mild cognitive impairment by volumetric and diffusivity measurements of the hippocampus. Psychiatry Res. Neuroimaging 146, 283–287 (2006).

57. Hett, K. et al. Patch-based DTI grading: Application to Alzheimer’s disease classification. In International Workshop on Patch-based Techniques in Medical Imaging, 76–83 (Springer, 2016).

58. Cui, Y. et al. Automated detection of amnestic mild cognitive impairment in community-dwelling elderly adults: a combined spatial atrophy and white matter alteration approach. Neuroimage 59, 1209–1217 (2012).

59. Li, M., Qin, Y., Gao, F., Zhu, W. & He, X. Discriminative analysis of multivariate features from structural mri and diffusion tensor images. Magn. resonance imaging 32, 1043–1051 (2014).

60. Jack, C. R. et al. The Alzheimer’s disease neuroimaging initiative (ADNI): MRI methods. J. magnetic resonance imaging 27, 685–691 (2008).

61. Jahanshad, N. et al. Diffusion tensor imaging in seven minutes: determining trade-offs between spatial and directional resolution. In Biomedical Imaging: From Nano to Macro, 2010 IEEE International Symposium on, 1161–1164 (IEEE, 2010).

62. Manjón, J. V. & Coupé, P. volbrain: An online MRI brain volumetry system. Front. neuroinformatics 10 (2016).

63. Manjón, J. V., Coupé, P., Martí-Bonmatí, L., Collins, D. L. & Robles, M. Adaptive non-local means denoising of MR images with spatially varying noise levels. J. Magn. Reson. Imaging 31, 192–203 (2010).

64. Avants, B. B. et al. A reproducible evaluation of ANTs similarity metric performance in brain image registration. Neuroimage 54, 2033–2044 (2011).

65. Tustison, N. J. et al. N4ITK: improved N3 bias correction. IEEE transactions on medical imaging 29, 1310–1320 (2010).

66. Romero, J. E., Coupe, P. & Manjon, J. V. Hips: A new hippocampus subfield segmentation method. NeuroImage 163, 286–295 (2017).

67. Romero, J. E., Coupé, P. & Manjón, J. V. High resolution hippocampus subfield segmentation using multispectral multiatlas patch-based label fusion. In International Workshop on Patch-based Techniques in Medical Imaging, 117–124 (Springer, 2016).

68. Coupé, P., Manjón, J. V., Chamberland, M., Descoteaux, M. & Hiba, B. Collaborative patch-based super-resolution for diffusion-weighted images. NeuroImage 83, 245–261 (2013).

69. Manjón, J. et al. Nice: non-local intracranial cavity extraction. Int. J. Biomed. Imaging (2014).

70. Manjón, J. V. et al. Diffusion weighted image denoising using overcomplete local pca. PloS one 8, e73021 (2013).

71. Basser, P. J., Mattiello, J. & LeBihan, D. Mr diffusion tensor spectroscopy and imaging. Biophys. journal 66, 259–267 (1994).

72. Garyfallidis, E. et al. Dipy, a library for the analysis of diffusion MRI data. Front. neuroinformatics 8, 8 (2014).

73. Tong, T. et al. A novel grading biomarker for the prediction of conversion from mild cognitive impairment to Alzheimer’s disease. IEEE Transactions on Biomed. Eng. 64, 155–165 (2017).

74. Barnes, C., Shechtman, E., Finkelstein, A. & Goldman, D. Patchmatch: A randomized correspondence algorithm for structural image editing. ACM Transactions on Graph. 28, 24 (2009).

75. Sutour, C., Deledalle, C.-A. & Aujol, J.-F. Adaptive regularization of the NL-means: Application to image and video denoising. IEEE Transactions on image processing 23, 3506–3521 (2014).

76. Whitwell, J. L., Crum, W. R., Watt, H. C. & Fox, N. C. Normalization of cerebral volumes by use of intracranial volume: implications for longitudinal quantitative MR imaging. Am. J. Neuroradiol. 22, 1483–1489 (2001).

77. Dukart, J., Schroeter, M. L., Mueller, K. & Alzheimer’s Disease Neuroimaging Initiative. Age correction in dementia– matching to a healthy brain. PloS one 6, e22193 (2011).

78. Giraud, R. et al. An optimized patchmatch for multi-scale and multi-feature label fusion. NeuroImage 124, 770–782 (2016).

79. Hochberg, Y. & Tamhane, A. Multiple comparison procedures (John Wiley, 1987).

80. Nir, T. M. et al. Diffusion weighted imaging-based maximum density path analysis and classification of alzheimer’s disease. Neurobiol. aging 36, S132–S140 (2015).

81. Maggipinto, T. et al. Dti measurements for alzheimer’s classification. Phys. Medicine Biol. 62, 2361 (2017).

82. Arbabshirani, M. R., Plis, S., Sui, J. & Calhoun, V. D. Single subject prediction of brain disorders in neuroimaging: Promises and pitfalls. NeuroImage 145, 137–165 (2017).

83. Thal, D. R. et al. Alzheimer-related *τ*-pathology in the perforant path target zone and in the hippocampal stratum oriens and radiatum correlates with onset and degree of dementia. Exp. neurology 163, 98–110 (2000).

84. Mueller, S. et al. Measurement of hippocampal subfields and age-related changes with high resolution MRI at 4T. Neurobiol. aging 28, 719–726 (2007).

85. Carlesimo, G. A. et al. Atrophy of presubiculum and subiculum is the earliest hippocampal anatomical marker of Alzheimer’s disease. Alzheimer’s & Dementia: Diagn. Assess. & Dis. Monit. 1, 24–32 (2015).

